# Proteome-wide prediction of bacterial carbohydrate-binding proteins as a tool for understanding commensal and pathogen colonisation of the vaginal microbiome

**DOI:** 10.1101/2020.09.10.291781

**Authors:** François Bonnardel, Stuart M. Haslam, Anne Dell, Ten Feizi, Yan Liu, Virginia Tajadura-Ortega, Yukie Akune, Lynne Sykes, Phillip R. Bennett, David A. MacIntyre, Frédérique Lisacek, Anne Imberty

**Affiliations:** University Grenoble Alpes, CNRS, CERMAV, Grenoble, France; Swiss Institute of Bioinformatics, Geneva, Switzerland; Computer Science Department, UniGe, Geneva, Switzerland; Department of Life Sciences, Imperial College London, London, UK; March of Dimes European Prematurity Research Centre, Imperial College London, UK; Glycosciences Laboratory, Department of Metabolism Digestion and Reproduction, Imperial College London, London, UK; Imperial College Parturition Research Group, Division of the Institute of Reproductive and Developmental Biology, Department of Metabolism Digestion and Reproduction, Imperial College London, London, UK; Queen Charlotte’s Hospital, Imperial College Healthcare NHS Trust, London, UK; Tommy’s National Centre for Miscarriage Research, Imperial College London, London, UK; Section of Biology, UniGe, Geneva, Switzerland

**Keywords:** Lectins, glycans, vaginal microbiota, pregnancy, infection, bioinformatics

## Abstract

Bacteria use protein receptors called lectins to anchor to specific host surface sugars. The role of lectins in the vaginal microbiome, and their involvement in reproductive tract pathophysiology is poorly defined. Here we establish a classification system based on taxonomy and protein 3D structure to identify 109 lectin classes. Hidden Markov Model (HMM) profiles for each class were used to search bacterial genomes, resulting in the prediction of >100 000 bacterial lectins available at unilectin.eu/bacteria. Genome screening of 90 isolates from 21 vaginal bacterial species showed that potential pathogens produce a larger variety of lectins than commensals indicating increased glycan-binding potential. Both the number of predicted bacterial lectins, and their specificities for carbohydrates correlated with pathogenicity. This study provides new insights into potential mechanisms of commensal and pathogen colonisation of the reproductive tract that underpin health and disease states.

## Introduction

Microbiota-host interactions within different ecological niches of the human body are critical determinants of health and disease states (Cho and Blaser, 2012). At mucosal surface interfaces, microbial and host cells, as well as non-cellular components of the mucosa, present an exceptionally complex array of attachment and recognition sites for microbiota, many of which are carbohydrate sequences displayed on extensively glycosylated mucin-type glycoproteins rich in O-glycans. The diverse populations of glycans provide recognition sites for microbial adhesins that have the ability to distinguish the various motifs displayed. Bacteria also produce glycosylhydrolases and other enzymes that facilitate the use of secreted mucins as primary carbon sources for energy metabolism (Tailford et al., 2015; Thornton et al., 2008). The abilities of microbes to specifically recognise, attach and adhere to cellular and non-cellular surfaces are thus key aspects of commensal and pathogenic colonisation and are mediated by receptors, such as lectins and carbohydrate-binding modules (CBMs) (Corfield, 2018; Etzold and Juge, 2014; Ficko-Blean and Boraston, 2012; Tailford et al., 2015).

Lectins are ubiquitous proteins of non-immune origin that bind to a variety of carbohydrates without modifying them (Lis and Sharon, 1998). Through their interactions with glycoproteins and glycolipids via the oligosaccharides, lectins play crucial roles in cell-cell communication, signalling pathways and immune responses (Lepenies and Lang, 2019). Bacterial lectins may be incorporated into multiprotein organelles, such as fimbriae (pili) or flagellae and participate in the mediation of host recognition and adhesion (Moonens and Remaut, 2017). In pathogenic species, lectins may also be subunits associated with a toxic catalytic unit that targets subcellular components (Merritt and Hol, 1995). Soluble lectins are also expressed as virulence factors by opportunistic bacteria (Imberty et al., 2005) and can alter dynamics of glycolipids to induce the internalization of whole bacteria into host cells (Eierhoff et al., 2014). Bacterial lectins have also been shown to directly impair immune signalling and repair pathways and are implicated in the formation of biofilms (Fazli et al., 2014).

The role of lectins and their ligands in shaping microbial niches in the human body is increasingly recognised, particularly at mucosal interfaces including the gut (Iliev et al., 2012; Pang et al., 2016; Tailford et al., 2015) and oral cavity (Cross and Ruhl, 2018). However, much less is known about the role of lectins in shaping microbial niches in the lower female reproductive tract, which play a key role in shaping health and disease throughout a woman’s life span (MacIntyre et al., 2017). Colonisation in the vagina by *Lactobacillus* species has for long been considered a hallmark of health (Ma et al., 2012; van de Wijgert et al., 2014), whereas *Lactobacillus* deplete, high diversity vaginal microbiomes enriched in potential pathogens are characteristic of bacterial vaginosis and are associated with increased risk of sexually transmitted infections (STIs) (Borgdorff et al., 2014; Reimers et al., 2016), progression of cervical cancer (Mitra et al., 2020) and adverse pregnancy outcomes such as miscarriage and preterm birth (Al-Memar et al., 2019; Brown et al., 2019; Fettweis et al., 2019; Kindinger et al., 2016). A key component of the vaginal mucosa are highly glycosylated mucins that are derived from the mucin-secreting glands of the cervix (Gipson, 2001). Alteration of terminal glycan residues of mucins by microbially secreted sialidases and sulphatases modulate the physical and immunological properties of the vaginal mucosa (Wiggins et al., 2001). Vaginal pathogens such as *Gardnerella vaginalis, Trichomonas vaginalis, Prevotella* and *Ureaplasma* species are capable of degrading secretory IgA (Cauci et al., 1998; Coombs and North, 1983; Kilian et al., 1996; Robertson et al., 1984). Moreover, specific strains of *Streptococcus agalactiae* (group B streptococci) secrete hyaluronidases that degrade cervical hyaluronic acid into disaccharide fragments dampening host immune activation through inhibition of Toll-like receptors, and thereby may contribute to preterm birth via ascending infection (Vornhagen et al., 2016). *Streptococcus agalactiae* can also implement a negative signalling mechanism known as sialoglycan mimicry to evade detection and phagocytosis by neutrophils; this is through terminal α2-3-linked sialic acids on the bacteria as ‘self’ glycans recognized by the neutrophil lectin Siglec-9 (Carlin et al., 2009).

Despite their important role in infection and pathogenicity, the full extent of the contribution of bacterial lectins to health and disease states is yet to be fully elucidated. This is partly due to the limited annotation and characterisation of the lectins in protein and proteome databases, that precludes predictions of the diversity, structure and function of the lectins. In recent years, this has begun to be addressed through the development of databases for structural and functional glycobiology (Mariethoz et al., 2018; Mariethoz et al., 2016). Among these, UniLectin3D provides 3D structures of more than 2500 lectins and their complexes with carbohydrates (Bonnardel et al., 2019) and sites within UniLectin, a platform dedicated to the curation and collection of lectin knowledge. In this study we describe how manual selection of lectin domains in 3D structures permits the identification of lectin classes characterised by fold similarity and minimum thresholds of sequence identity. We show that defined amino acid sequence motifs and profiles characterising each lectin class can be used to screen proteomes and translated genomes to identify unannotated lectins. Comparison of these lectins across different vaginal microbiota strains provides new insights into the potential mechanisms by which commensal and pathogen colonisation are associated with physiological and pathological conditions in the lower reproductive tract.

## Results

### Structural classification of bacterial lectins in Unilectin3D

Structural classification of known lectins in the curated Unilectin3D database (www.unilectin.eu/unilectin3D/) was performed first on the basis of differences in fold, i.e. structure of the protein backbone and second on amino acid sequences at 20% of sequence similarity for lectin classes and at 70% of sequence similarity for lectin families. This led to the identification of 35 different folds and 109 lectin classes (S1 Table) derived from a total of 2483 structural lectin entries that primarily originated from plant and animal sources. However, bacterial lectins from 46 different species accounted for approximately 20% of database entries (495/2483), which were distributed among 19 different folds (Figure 1) and 37 classes (S1 Table).

**Figure 1:**
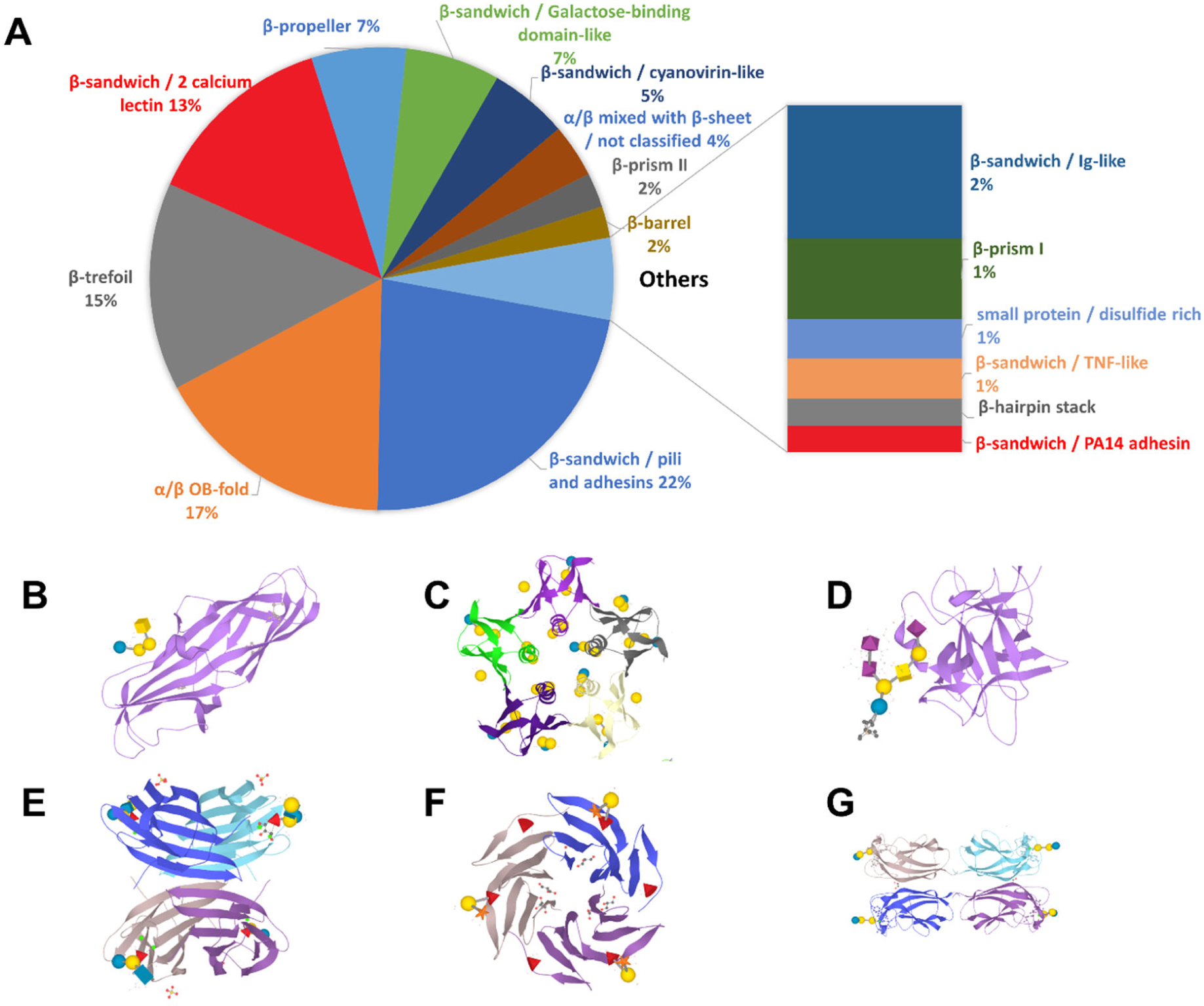
Structural classification of bacterial lectins. (A) Distribution of bacterial lectin folds derived from the UniLectin3D database. From the analysis of fold distribution of bacterial lectin crystal structures, the six most frequent fold are represented: (B) Pili and adhesins: 1J8R PapG *Escherichia coli*, (C) OB fold: 1BOS SLT-1 / STX-1 *E. coli*, (D) *β*-trefoil: 1FV2 TeNT *Clostridium tetani*, (E) 2 calcium lectin: 1W8F LecB / PA-IIL, RSIIL *Pseudomonas aeruginosa*, (F) *β*-propeller: 2BS6 RSL, BambL *Ralstonia solanacearum*, and (G) Galactose binding domain-like: 2VXJ LecA / PA-IL *Pseudomonas aeruginosa*. 3D structures were generated using LiteMol (Sehnal et al., 2017) with monosaccharides in binding sites represented using Symbol Nomenclature for Glycans (SNFG) (Neelamegham et al., 2019).

The analysis of fold distribution in bacterial lectin crystal structures showed an over-representation of β-sheet containing folds, which were common to adhesins and toxins including previously described pili adhesins, such as FimH in uro-pathogenic *Escherichia coli*, the oligomer-binding (OB) fold of the cholera toxin binding domain, the β-sandwich of LecA and LecB in *Pseudomonas aerigunosa* and the β-trefoil of the recognition domain in clostridial neurotoxins. While the majority of folds were shared between lectins of different origins, classes of pili adhesins and AB5 toxins were found to be restricted to bacteria.

### Prediction of lectin sequences from bacterial proteomes

Alignment of amino acid sequences in each of the 109 identified lectin classes led to the identification of 109 characteristic motifs of conserved residues. Profiles characterising each lectin class were generated with Hidden Markov Models (HMM), which were subsequently used to screen 130 million bacterial protein sequences from the UniProt database and over 168 million bacterial protein sequences from the NCBI RefSeq database derived from over 100 000 bacterial species. The TIM fold (named after triosephosphate isomerase) and Variable Lymphocyte Receptor folds are highly frequent in the resulting predictions. The TIM lectin class may arise from its high occurrence in hydrolases. Consequently, both of these lectin classes were excluded from whole proteome predictions. This resulted in the selection of 100 671 sequences as putative lectins in 10126 distinct bacterial species (reduced to 46 322 sequences in 6 425 distinct bacterial species when applying a score of 0.25). A web interface dedicated to the exploration of these bacterial lectin candidates is available at www.unilectin.eu/bacteria/.

Although 481 3D-structures of bacterial lectins are categorized into 37 classes, the screening results indicated that putative bacterial lectins occur in 97 out of the 107 identified classes (with a cutoff of 25% of sequence similarity with the reference) (S1 Table). Putative lectin sequences identified in each class, together with the distribution of the prediction scores to the original HMM motif, are presented in Figure 2. The distribution of the fold predicted for bacterial lectins differs from that obtained when using 3D structures generated from the UniLectin3D database with several classes comparatively over-represented, including the Ricin-like (*β* trefoil), the LysM domains (LysM fold), and the F-type lectins (*β* sandwich galactose binding domain like) (S1 Table). Each lectin domain is predicted by selecting the best fitted HMM. A score reflecting the sequence similarity is computed as the difference between the predicted lectin domain and the reference conserved motif. Lectins with the highest prediction scores per class were, as expected, of bacterial origin and included adhesins, AB5 toxins and calcium-dependent soluble lectins. However, the β-prism III fungal lectin was also found to have a high prediction score indicative of genetic exchange between bacteria and fungi. The majority of low scoring predictions (<0.25) reflective of low sequence similarity, were those associated with viruses with the exception of the Influenza hemagglutinin, which contains a high abundance of sequences for the characteristic domain although not all are carbohydrate-binding. Lectins with mid-range (0.25-0.5) prediction scores were evenly distributed across multiple genome sources.

**Figure 2:**
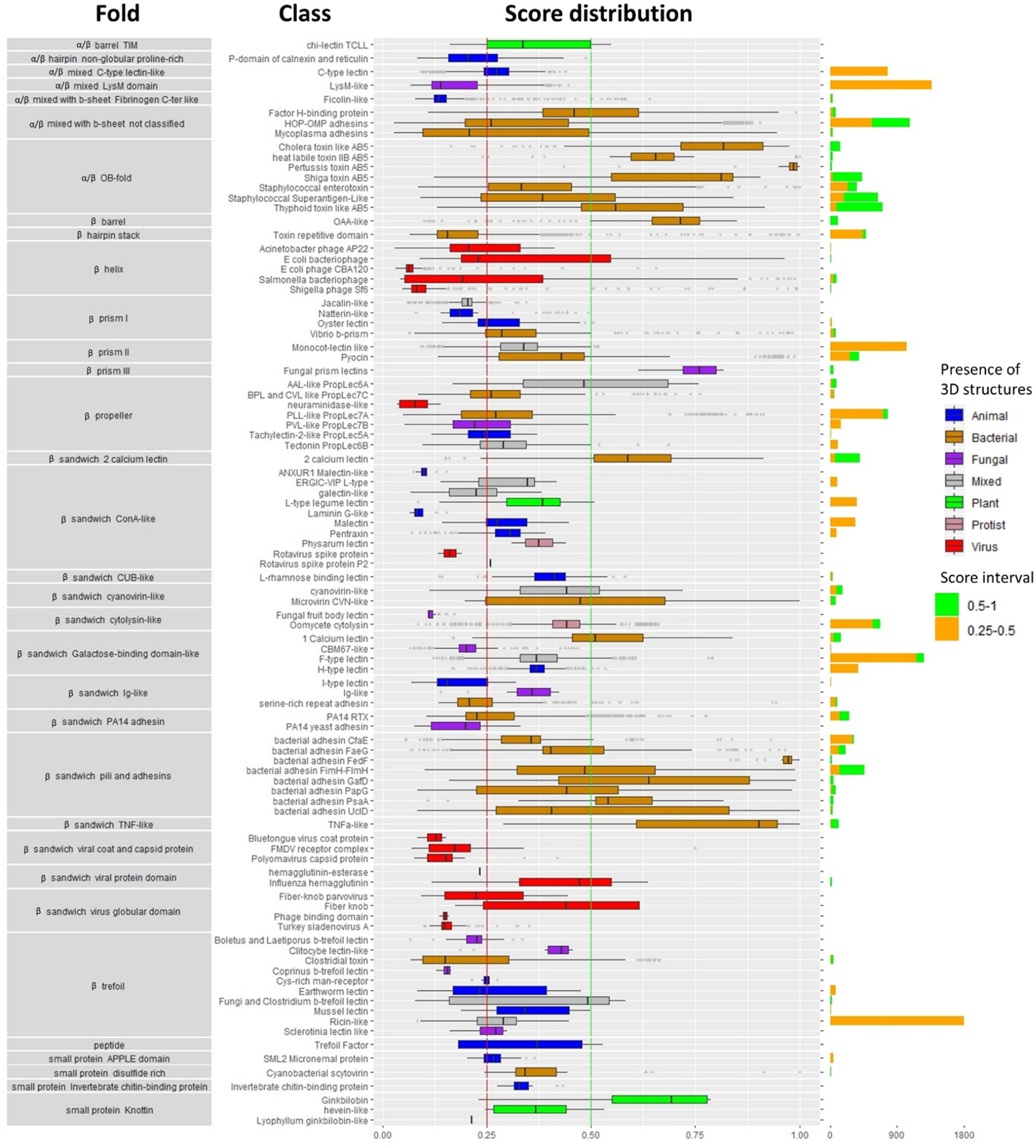
Distribution of structural fold-types within predicted lectin classes derived from 21 different bacterial genomes. Distributions of the predicted lectin classes are presented as horizontal box and whisker plots coloured on the basis of genome origin. The whisker plot represents the minimum, maximum, median, first quartile and third quartile in each class. Values approaching 1 are indicative of high sequence similarity to the reference motif. The predicted lectins in [0.25-0.5] and [0.5-1] score intervals are presented as bar graphs. The total number of predicted lectins in each class is listed in S1 Table.

### Identification and characterisation of vaginal microbiota lectins

We next obtained publicly available genome data for 90 vaginal bacterial strains classified on the basis of potential pathogenicity within the vaginal niche and having a known association with states of health or disease (S2 Table). Comparison of the lectomes, i.e. the predicted ensemble of lectins, highlighted major differences across species with pathobionts generally harbouring a higher diversity of lectin classes compared to commensals (Figure 3). Considering the low number of identified lectins, the TIM lectin and the Variable lymphocyte receptor classes were kept, despite a low probability of lectin activity. For example, the only predicted lectin consistently identified across *L. crispatus* isolates is LysM, a common domain involved in cell wall attachment in many different bacteria. Consistent with this, the LysM domain was predicted in the majority of vaginal microbial genomes examined but interestingly, was absent from *L. iners* and most *Provotella* strains.

**Figure 3.**
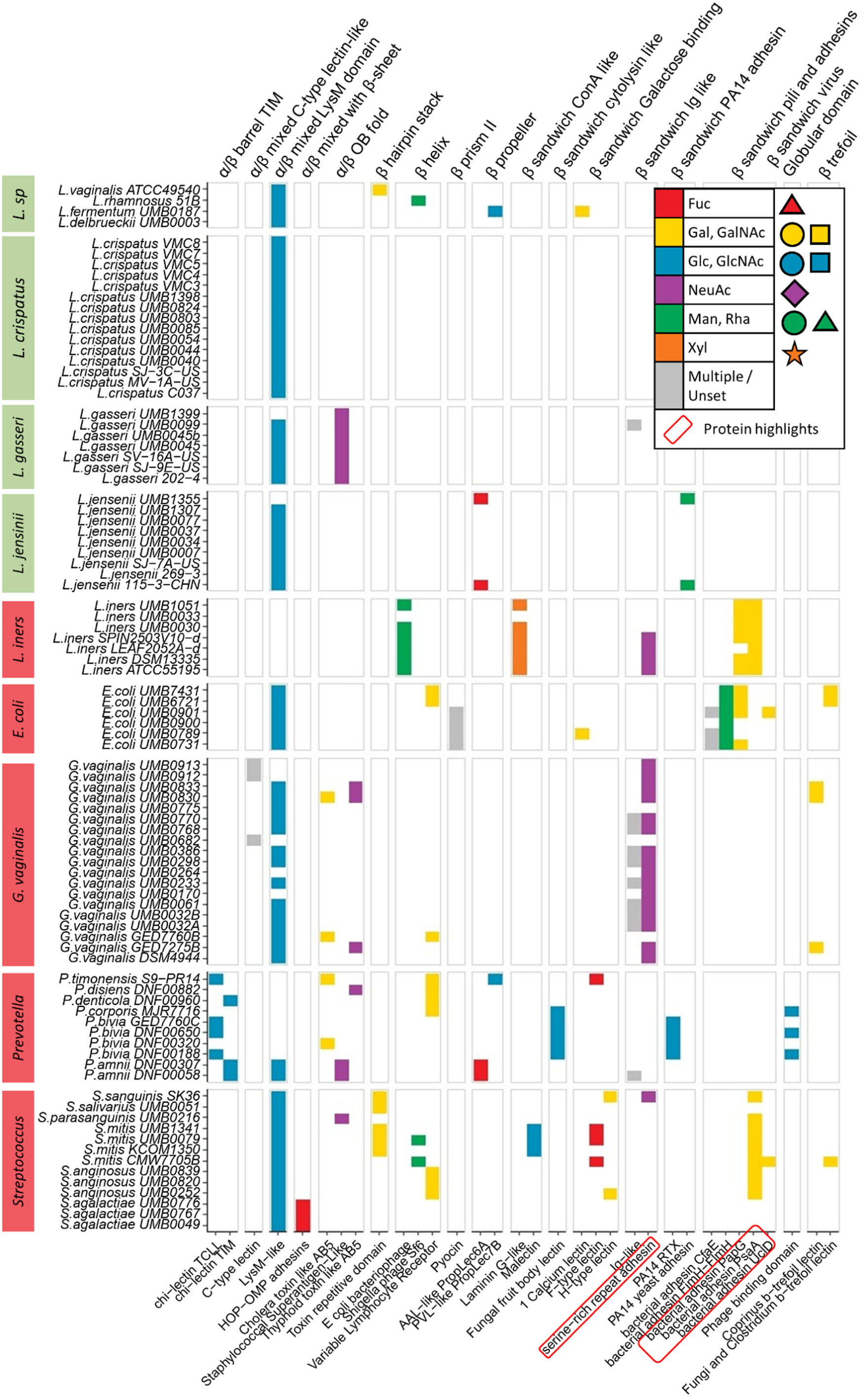
Heatmap of predicted lectomes from different vaginal commensal and pathobiont bacterial species classified by fold and class. Green species label represent commensal species and red species labels represent pathobiont species. Colours within each class of lectin reflects its main glycan specificity characterised by binding monosaccharides using standardised Symbol Nomenclature for Glycans (SNFG) (https://www.ncbi.nlm.nih.gov/glycans/snfg.html). The lectin class circled in red are further discussed in the results due to their particular presence in *L*.*iners*.

Predicted lectins of *L. iners* could be mapped to five different classes: *E*.*coli* bacteriophage β-helix, laminin G-like, adhesin domain of two type 1 pili PapG and PsaA (chaperon-usher-assembled, CUP) and the adhesin domain of serine rich repeat protein (SRRP), which was also observed prominently in *G. vaginalis* species and in a *Streptococcus sanguinis* strain. Up to 10 different lectin classes were predicted from other *Streptococcus* species although *S. agalactiae* (also known as Group B Streptococcus), which is a pathogen known to cause sepsis, pneumonia and meningitis in newborn babies, was the only vaginal species predicted to produce Outer Membrane Protein (OMP) adhesins.

### Predictions of carbohydrate binding modules in vaginal microbiota

The screening strategy was extended to the prediction of carbohydrate binding modules (CBMs), which occur as small domains generally associated with carbohydrate modifying enzymes, often involved in microbial digestion of mucin glycans. A few of particular interest including CBM34, CBM41 and CBM48, which are specific for glucose containing polysaccharides (e.g. amylose, glycogen), these generally act as binding modules for amylases and related enzymes, and are predicted consistently across almost all vaginal species (Figure 4). While the majority of CBMs have been characterised as enzyme-associated domains in plant polysaccharides, two human-specific CBMs were observed in the dataset (S3 Table). The first is CBM40 considered as sialic acid-specific as it has been identified in association with a bacterial sialidase (Boraston et al., 2007). In the dataset analysed here, it is predicted to occur only in *L. iners* pathobiont species, *S. mitis* and some *Prevotella* species. Considering the earlier observation regarding the predicted SRR adhesin domain, the sialic acid binding ability appears to correlate mainly with lectins and CBMs present in the lectomes of pathobiont bacteria. The second domain of interest is CBM47, shown to be fucose-specific in the lectin regulatory domain of a cholesterol-dependent cytolysin present in some *S. mitis* strains (Feil, Lawrence et al. 2012). It shares structure and sequence similarity with the F-lectins from fishes (Vasta et al., 2017). In our study, this fucose-binding module is identified in *S. mitis* as well as in some pathobionts, i.e. *Provotella* and *L. iners*. Furthermore, the *L. iners* lectome contained two predicted adhesins, PapG and PsaA, as well as CBM60, which bind to galactose epitope occurring on human Gb3 gangliosides (Montanier et al., 2010).

**Figure 4.**
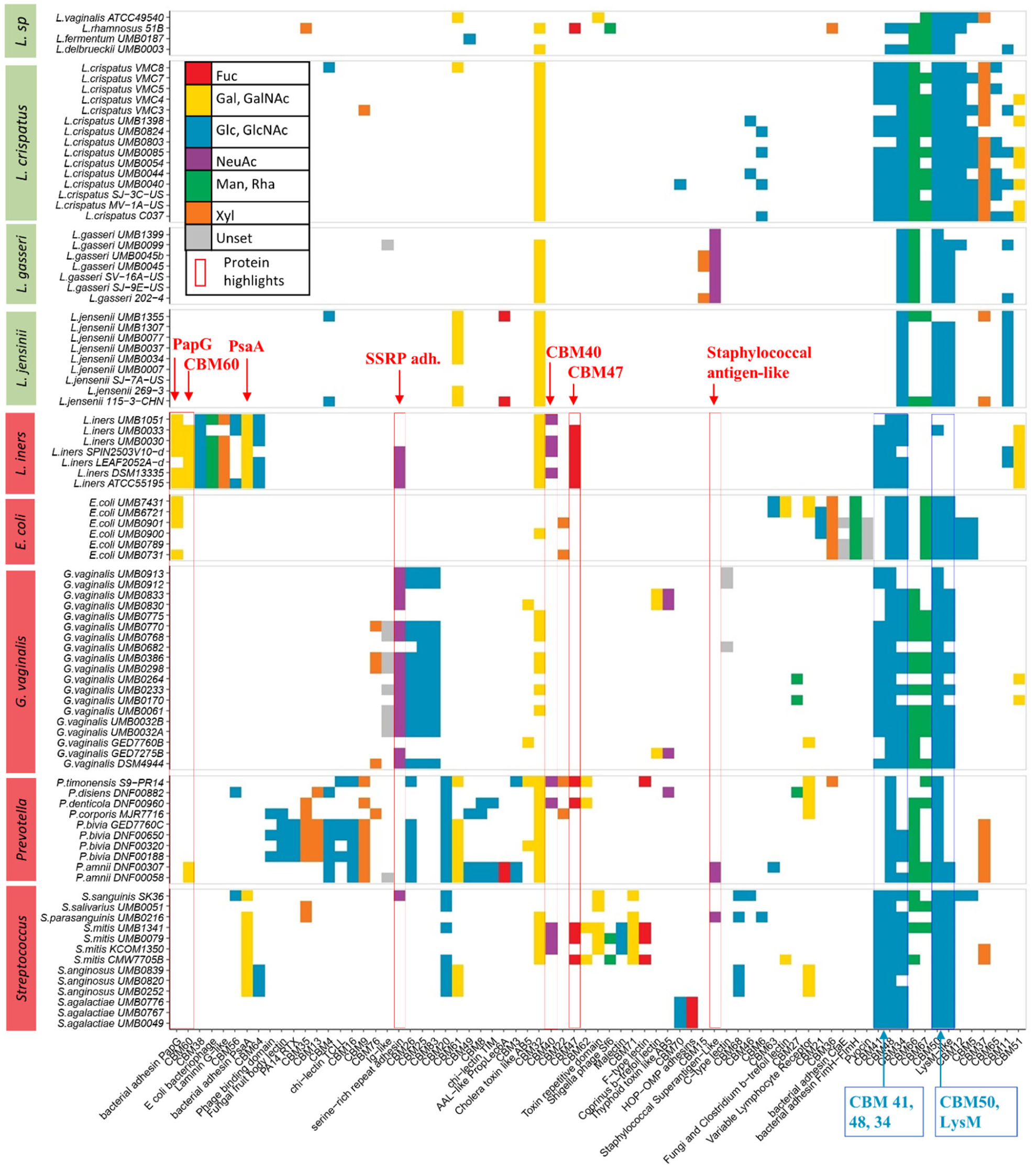
Heatmap of predicted lectin and CBM domains from different vaginal commensal and pathobiont bacterial species arranged by domain composition similarity. Colours within each class of lectin reflect its main glycan specificity (SNFG nomenclature) referred to in Figure 3. The domains highlighted are further discussed in the results due to their presence in *L. iners*. The inclusion of the CBM domains strengthens the distinction between commensal and pathobiont bacteria.

Further comparison of the predicted lectin and CBM profiles of vaginal commensals and pathobionts was made by performing unsupervised hierarchical clustering on a Euclidean distance matrix of the number of proteins per species for each lectin and CBM domain (Figure 5). The resulting hierarchical radial plot using predicted lectins showed a clear clustering of the majority of *Lactobacilli*, with further sub-clustering at species level observable. *L. iners* strains were an exception as they clustered more closely with other pathobiont species including *Prevotella* and *Streptococcus* species (Figure 5A). *G. vaginalis* also did not cluster in a single group. The inclusion of predicted CBMs in the clustering led to improved discrimination between commensal and pathobiont species and led to species-specific clustering of the majority of isolates (Figure 5B).

**Figure 5.**
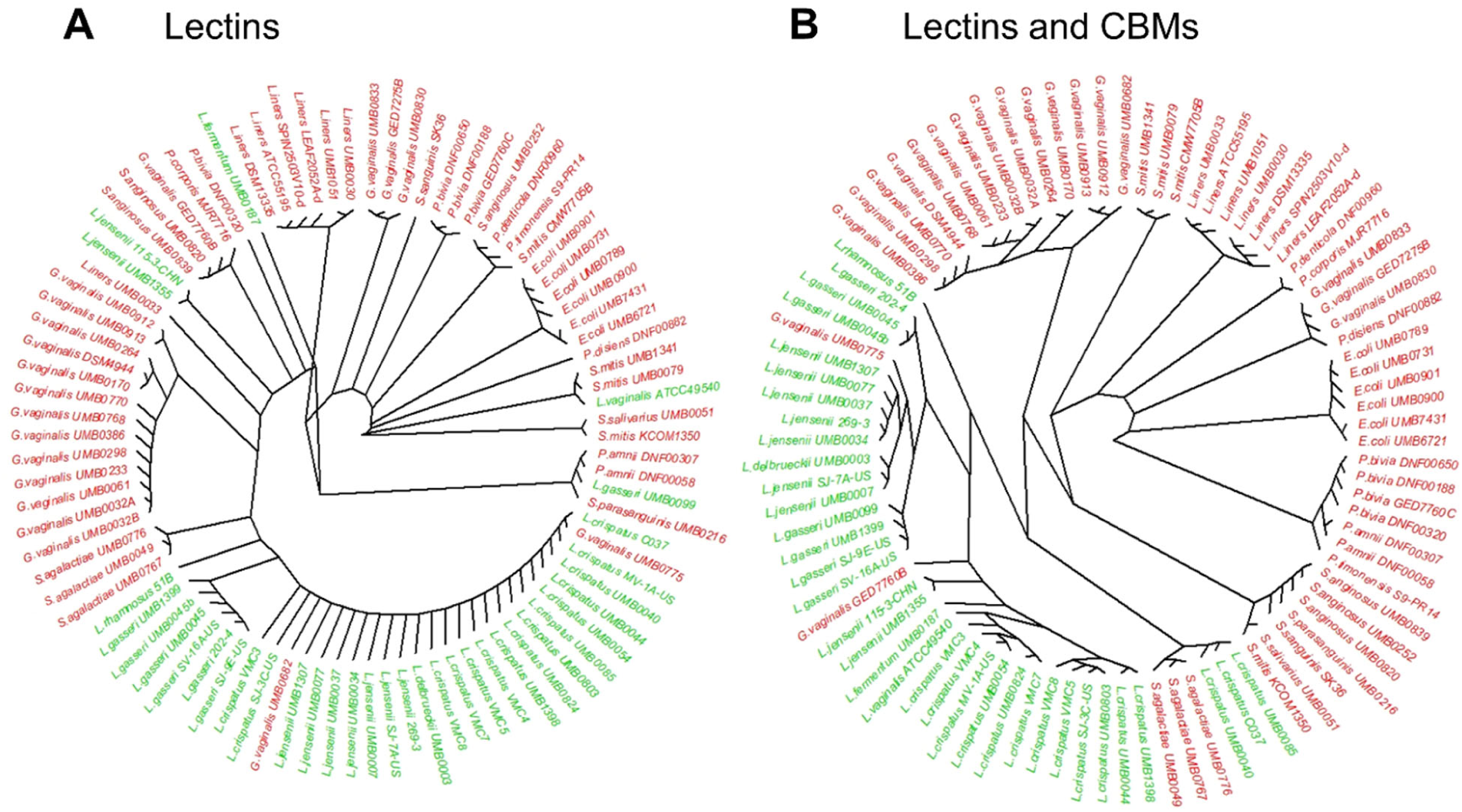
Hierarchical radial tree of (A) predicted lectin classes only or (B) lectin classes and predicted CBMs in vaginal commensal (green) and pathobiont (red) bacteria. LysM and CBM50 are excluded from the dataset to generate the hierarchical radial tree. While the majority of *Lactobacillus* species clustered closely to each other, indicating similar putative lectomes, the lectome of *L*.*iners* isolates more closely resembled that of pathobionts

## Discussion

The contribution of bacterial lectins to health and disease remains poorly understood. This is in part because their structural and functional complexity and the limited annotation of bacterial lectins in protein and proteome databases has prevented the development of predictive models of structure, diversity and function. Here, we begin to address this through manual selection of lectin domains in 3D structures obtained from the recently curated Unilectin3D database, followed by the prediction of lectin classes based upon fold similarity and minimum thresholds of sequence identity. This strategy has led to the identification of more than 35 different structural folds and 109 predicted lectin classes, of which 19 folds and 37 classes were of bacterial origin. These were particularly rich in β-sheet containing folds, which have previously been recognised as key structural characteristics of lectins from non-bacterial origin (Loris, 2002). Moreover, predicted classes of pili adhesins and AB5 toxins were found to be exclusive to bacteria. While other lectin classes also appeared to be exclusively predicted in bacteria, these results are likely to be influenced by the fact that to date, many structurally characterised and curated lectins represent those of highest abundance in readily culturable bacteria.

The predicted lectins in the bacterial proteomes and their specificities of glycan binding promise to be a basis for future designs of therapeutic molecules to target specific pathogenic bacteria. Detailed knowledge on the glycan binding specificities of the proteins is eagerly awaited. Glycan array analyses of the whole bacterial cells (now in progress) and of the recombinantly expressed proteins and determination of their 3D structures are envisaged to elucidate and validate the predicted recognition of glycans.

Given the increased awareness of the importance of the vaginal microbiome in shaping reproductive tract health outcomes, we next compared the occurrence of lectin-like proteins among vaginal commensal and pathogenic bacteria isolated from the vagina. Our analysis, based on 109 structurally characterised lectin classes, suggests that the common commensal species, *L. crispatus* and *L. gasseri*, produce only LysM, which is a ubiquitous domain detected in almost all bacteria. Previous studies of this domain have shown that it can bind peptidoglycan with a specificity for N-acetylglucosamine (Mesnage et al., 2014). CBM50 is one of the other naming of LysM and is therefore also widespread. In the CAZy database CBM50 is annotated as being involved in binding to N-acetylglucosamine residues in bacterial peptidoglycans and chitin The number of CBMs identified in the bacterial species investigated in the present study is small; these correspond mainly to domains associated with nutrient-degrading glycosylhydrolases. These results suggest that *Lactobacillus* species associated with optimal vaginal microbiome compositions appear to be comparatively ill-equipped for binding mucins. It is important to note that this observation may be biased because the analysis only involved structurally characterised lectins. Further, a limited number of other “mucin adhesion factors’’ have been described in *Lactobacilli* (Nishiyama et al., 2016; Velez et al., 2007), but except for the fimbriae domain in *L. rhamnosus* (Nishiyama, Ueno et al. 2016), these are in general described as moonlighting proteins, i.e. with adhesion properties being only a side activity in addition to their main function.

A shift from *Lactobacillus* species dominance of the vaginal niche towards increased bacterial diversity and enrichment of pathobionts is a signature of vaginal dysbiosis, which has been associated with a range of pathology states including increased risk of sexually transmitted infections (Borgdorff et al., 2014) and various poor pregnancy outcomes including miscarriage (Al-Memar et al., 2019), prelabour premature rupture of the foetal membranes (Brown et al., 2019; Brown et al., 2018) and preterm birth (Fettweis et al., 2019; Kindinger et al., 2017; Kindinger et al., 2016; Vaneechoutte, 2017). We propose that the strategy for binding glycans has evolved more in vaginal pathobionts than in commensals, with the former producing a much larger variety of lectins and CBMs. Our findings are consistent with previous reports on the occurrence of a large number of lectin domains in different species of *Streptococci;* these participate in the architecture of toxins, adhesins and pilins (Moschioni et al., 2010).

Whereas *Lactobacillus* species are considered hallmarks of optimal vaginal health, *L. iners* is considered a marker of a “transitional microbiome” at the crossroads of vaginal health and disease (Macklaim et al., 2011; Petrova et al., 2017). The predicted lectomes of the various *L. iners* strains screened were found to contain a significantly larger number of lectin-like domains than those in other *Lactobacilli*, and the same applies to our analyses of CBM-like domains. This is somewhat surprising considering that *L. iners* genome is much smaller than those of other *Lactobacilli* (Macklaim et al., 2011). Several of these identified domains are resemble proteins that are able to bind to glycans present on human mucins, examples are sialic-binding domain from SRPPs, galactose-specific pilin domain, as well as fucose-binding CBMs usually associated with *Streptococci*. This similarity between *L. iners* and pathogens is in agreement with the previous identification of inerolysin, a pore-forming toxin from *L. iners* also found in *Gardnerella* (Rampersaud et al., 2011). Moreover, sequences with similarity to fimbrial proteins PapG from *E. coli*, and Psa/Myf from *Yersinia pestis* were identified in almost all strains of *L. iners*. Interestingly, these two adhesins have similar specificity towards α-galactosylated epitopes (Kline et al., 2009).

The lectome expansion that appears to correlate with the transition towards species involved in vaginal dysbiosis, raises the question of concomitant changes in vaginal glycans, perhaps in glyco-epitopes present on mucins. Mucin glycans have been well investigated in gut and lung, and it has been demonstrated that glycosylation is altered in case of inflammation. For example, in cystic fibrosis patients, inflammation results in an increase in fucosylation and sialylation, favouring the attachment of opportunistic pathogens such as *Pseudomonas aeruginosa*, which in turn stimulates the inflammatory process (Cott et al., 2016). Such glycan-based processes may occur in the vagina and a deeper characterisation of mucin glycosylation in this context is needed.

While the mechanisms underpinning dynamic shifts in vaginal microbial structure and composition remain to be fully elucidated, our study provides important new insights into lectin profiles of commensal and pathogen colonisation of the reproductive tract that are associated with health and disease states.

The bioinformatics screening tools described and used in the present study can be run on any protein sequence data to reveal information currently lacking on the content and the role of the lectome. Results show clearly the emergence of characteristic patterns indicative of pathological states. This may guide the development of new strategies for novel therapeutics designed to manipulate adhesion and attachment of microbes to promote optimal colonisation of the lower reproductive tract.

## Acknowledgments

The authors acknowledge support by the ANR PIA Glyco@Alps (ANR-15-IDEX-02), Alliance Campus Rhodanien Co-Funds (http://campusrhodanien.unige-cofunds.ch), Labex Arcane/CBH-EUR-GS (ANR-17-EURE-0003) and the March of Dimes.

## Author Contributions

A. I., F. L. and D. A. M. developed the concept, supervised the research and wrote the original draft. F.B. designed the database and conducted the research. S.M.H., A.D., T.F. and Y.L. validated the observation and participated in the redaction. V. T.-O., Y.A., L.S., P.R.B. participated in the analysis of data and in the revision of manuscript.

## STAR Methods

***Key resources table***

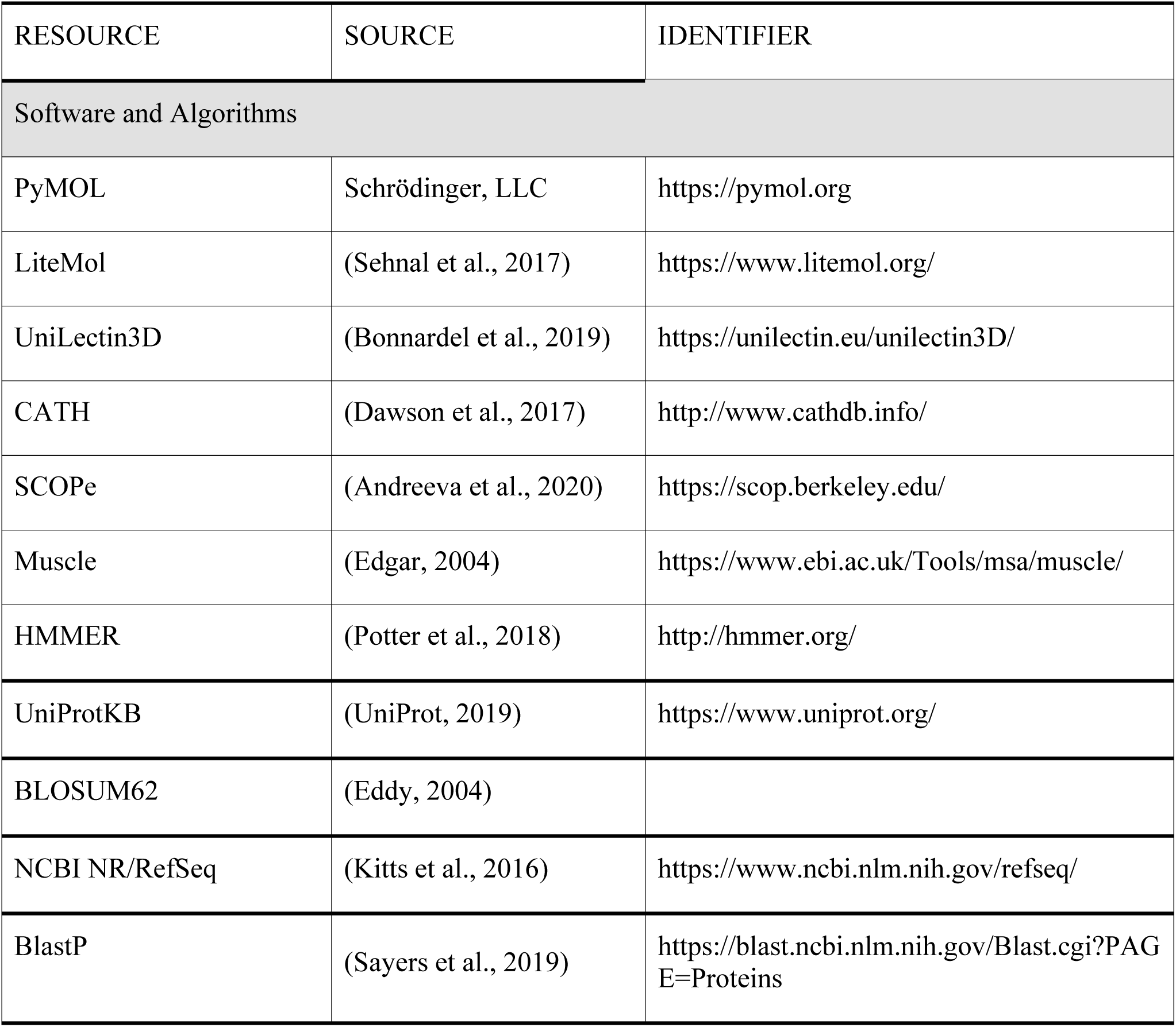
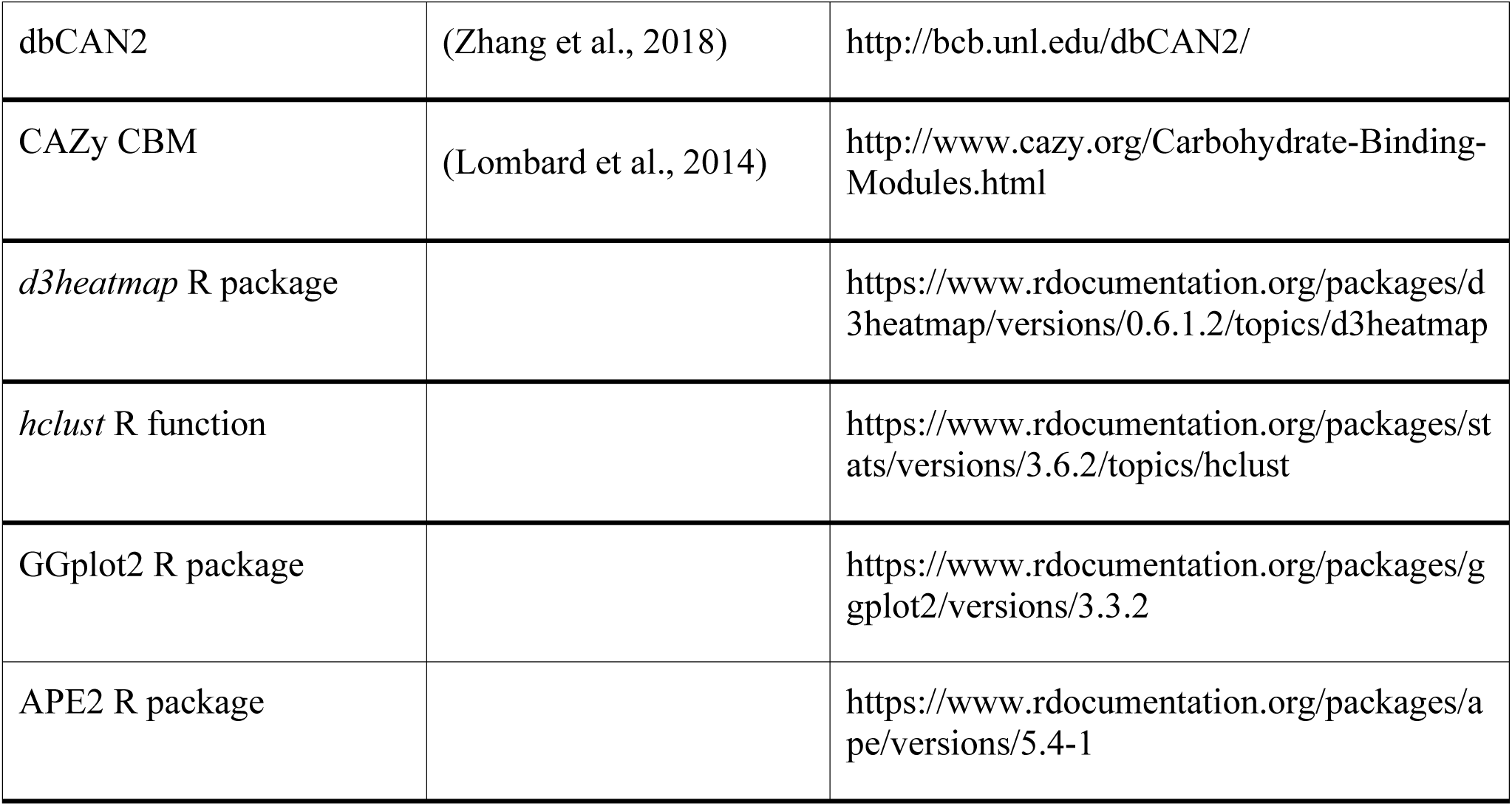

### Definition of signature profiles for lectins

A new lectin classification has been recently defined based on structural data and is available in the UniLectin3D database (https://unilectin.eu/unilectin3D/). The classification is built on three levels: 1) the fold level directly derived from the protein three-dimensional structure that describes the fold adopted by the whole lectin domain (β-helix, β-propeller and others). The nomenclature on fold are adopted from the reference structural-based databases, CATH (Dawson et al., 2017) and SCOPe (Andreeva et al., 2020) and previous reports on structural classification of lectins (Fujimoto et al., 2014); 2) The class level defined by sequence similarity with a 20% cut-off between different classes, i.e., lectin sequences in one class are at least 20% similar to one another; 3) The family level defined at a minimum of 70% of sequence identity. The values of cut-offs were set in agreement with definitions in the CATH database for the class level, and empirically for the family level in order to maximise the consistency of each family. The classification is therefore organized in 35 folds, 109 classes, and 350 families.

For each of the 109 lectin classes, UniLectin3D sequences were aligned with the Muscle software (Edgar, 2004) to construct a characteristic motif of conserved residues. Sequence redundancy was automatically removed. Manual inspection of characteristic lectin domains led to creating a list of disqualifying domains such as peptide tags in order to manage future systematic removal. Conserved regions from the multiple alignments were then fed to a Hidden Markov Modelling tool to generate profiles characterising each lectin class. The HMMER-hmmbuild tool (Potter et al., 2018) was used to align each lectin class multiple sequence alignment against protein sequence datasets, with the sym_frac parameter at 0.8 to avoid isolated regions in the conserved motifs.

### Prediction of bacterial lectins in protein databases

Bacterial sequences recorded in UniProtKB (UniProt, 2019) and in non-redundant NCBI were processed with HMMER-hmmsearch, with default parameters and a p-value below 10^-2^, to run profiles obtained with HMMER-hmmbuild. Parameters include the BLOSUM62 score matrix for amino acid substitutions (Eddy, 2004). Further filtering was applied to multiple strains of the same species with almost identical proteins and only a few different amino acids due to natural mutation, sequencing errors, or protein prediction errors. Post-processing involved keeping only one representative protein for all redundant proteins (with 100 consecutive amino acids that are identical). Predicted domains with less than 15 amino acids are considered as small fragments.

Each sequence match output by the HMMER toolset is evaluated with a quality score that has no upper boundary. Furthermore, because each family profile is generated independently of one another, quality scores are not comparable across motifs used for the prediction. This makes it impossible to use a single cut-off for all lectin classes. Additionally, in the case of tandem repeat domains, the quality score is proportional to the number of repeats and artificially promotes sequences with repeated domains. To address these scoring issues, a prediction score for each database hit was defined to give the similarity between the predicted domain and the reference lectin motif. The amino acid sequence alignment generated by HMMER during the search is further evaluated: at each position of the alignment, a cumulative counter is incremented by 1 if amino acids are identical, else by a normalised BLOSUM62 substitution score. The final value of the counter divided by the domain length (i.e., the total number of positions) results in a value between 0 to 1 that defines the prediction/similarity score. A predicted lectin may belong to several classes, independently of the prediction score. The prediction/similarity score is mainly destined to order the information to be displayed on the UniLectin platform for each predicted lectin. HMMER p-value threshold (better defined then HMMER score) applied before remains the most reliable parameter for trusting a candidate lectin.

For each predicted protein, associated annotations are extracted and loaded from UniProt and from the NCBI. This includes the taxonomy details of the protein and the corresponding ID of the NCBI taxonomy database. Proteins considered as obsolete in the latest releases of UniProt or in the NCBI, with no associated metadata, are removed.

### Prediction of lectins and CBMs in the vaginal microbiome

The subset of bacteria corresponding to the vaginal microbiome (S2Table) was identified from genome database annotations, such as those found in the Bioproject www.ncbi.nlm.nih.gov/bioproject/PRJNA316969 and from a published list of bacteria (Thomas-White et al., 2018). Bacteria belonging to different species of *Lactobacilli, Gardnerella, Prevotella, E. coli* and Group B *Streptococc*i were selected and classified into commensals or pathobionts on the basis of their potential pathogenicity within the vaginal niche (van de Wijgert et al., 2020), and their association with states of health and disease including bacterial vaginosis, preterm birth and risk of acquisition of sexually transmitted infections (Borgdorff et al., 2014; Brown et al., 2018; Fettweis et al., 2019; Kindinger et al., 2017; Ma et al., 2012; Petrova et al., 2017; van de Wijgert et al., 2020; van de Wijgert et al., 2014).

The proteome of each strain was downloaded from the NCBI assembly database (Kitts et al., 2016). The corresponding sequences were processed to detect lectins and CBMs with the same method of prediction involving the 109 lectin profiles generated as described above. HMMER-hmmsearch was run to identify the lectome of each strain’s proteome with default parameters and a p-value below 10^-2^ with no further filtering. Proteins producing good quality alignments (HMM score > 50) with HMMER during the analysis of amino acid sequences were directly tagged as lectin domains. For lesser quality alignments the “Align Sequences Protein BLAST” component of the BlastP tool (Sayers et al., 2019) was used with default parameters to align a predicted domain against the closest reference lectin with a defined 3D structure. Manual quality checks, especially focused on the glycan binding pocket, were carried out to verify the amino acid conservation and ensure the quality of the predicted lectin.

HMM profiles of Carbohydrate-binding modules (CBMs)were extracted from dbCAN2, a web server for the identification of carbohydrate-active enzymes (Zhang et al., 2018). The HMM profiles provided by dbCAN2 are based on CAZy CBM sequence data (Lombard et al., 2014). These profiles were used to identify 1777 proteins from the predicted proteomes of the vaginal commensals and pathobionts. Following removal of high frequency influenza-like predicted lectins and CBD domains occurring in less than three strains, the resulting data was grouped by domain clustering to reflect compositional similarities. The remaining CBMs were associated with their matching glycans and additional information (S3 Table).

To reinforce the results influenza-like predicted lectins are removed (the high frequency of this domain is misleading, as mentioned earlier) and the lectin and CBM domains occurring in less than three strains were filtered out (removing 20 lectin classes and 15 CBM domains for a total of 50 proteins).

### Prediction of lectins and CBMs in the vaginal microbiome

Predicted lectins in the HMMER output format were formatted into a tabulated matrix flat file by a python parser and loaded in R for statistical analysis. The following libraries were used:

1. Graphics were generated with R libraries of the Comprehensive R Archive Network (CRAN) including the *d3heatmap* package for heatmaps
2. Hierarchical clustering: The Ward’s minimum variance method part of the *hclust* R package was used to process a Euclidean distance matrix of the number of predicted proteins per species for each domain
3. GGplot2 and the APE (Analyses of Phylogenetics and Evolution) package for the hierarchical tree. In this case, prior clustering was applied to the data with the complete linkage method of the *hclust* R package. A Euclidean distance matrix of the number of predicted proteins per species for each domain was input.

For the sake of simplicity, lectins occurring in at least two strains are represented and the Influenza domain is filtered out for the lectin heatmap; and in at least 3 strains for the lectin and CBM heatmap. When lectins and CBMs are represented together the domains present in at least three strains are considered. The lectin and CBM specificity for glycans was manually recovered using UniLectin3D database and CAZy database annotations. Only predicted bacterial lectins with a score greater than 0.25 are kept.

## Supplemental Information

**S1 Table** : List of lectin classes identified from Unilectin3D and used in the classification

**S2 Table**. List of the species and strains used in the study

**S3 Table**. CBMs of interest for the present study with associated glycan specificity

## References

Al-Memar, M., Bobdiwala, S., Fourie, H., Manino, R., Lee, Y.S., Smith, A., Marchesi, J.R., Timmerman, D., Bourne, T., Bennett, P.R., et al. (2019). The association between vaginal bacterial composition and miscarriage: a nested case-control study. BJOG.

Andreeva, A., Kulesha, E., Gough, J., and Murzin, A.G. (2020). The SCOP database in 2020: expanded classification of representative family and superfamily domains of known protein structures. Nucleic Acids Res 48, D376–D382.

Bonnardel, F., Mariethoz, J., Salentin, S., Robin, X., Schroeder, M., Perez, S., Lisacek, F., and Imberty, A. (2019). UniLectin3D, a database of carbohydrate binding proteins with curated information on 3D structures and interacting ligands. Nucleic Acids Research 47, D1236–D1244.

Boraston, A.B., Ficko-Blean, E., and Healey, M. (2007). Carbohydrate recognition by a large sialidase toxin from Clostridium perfringens. Biochemistry 46, 11352–11360.

Borgdorff, H., Tsivtsivadze, E., Verhelst, R., Marzorati, M., Jurriaans, S., Ndayisaba, G.F., Schuren, F.H., and van de Wijgert, J.H. (2014). Lactobacillus-dominated cervicovaginal microbiota associated with reduced HIV/STI prevalence and genital HIV viral load in African women. ISME J 8, 1781–1793.

Brown, R.G., Al-Memar, M., Marchesi, J.R., Lee, Y.S., Smith, A., Chan, D., Lewis, H., Kindinger, L., Terzidou, V., Bourne, T., et al. (2019). Establishment of vaginal microbiota composition in early pregnancy and its association with subsequent preterm prelabor rupture of the fetal membranes. Transl Res 207, 30–43.

Brown, R.G., Marchesi, J.R., Lee, Y.S., Smith, A., Lehne, B., Kindinger, L.M., Terzidou, V., Holmes, E., Nicholson, J.K., Bennett, P.R., et al. (2018). Vaginal dysbiosis increases risk of preterm fetal membrane rupture, neonatal sepsis and is exacerbated by erythromycin. BMC Med 16, 9.

Carlin, A.F., Uchiyama, S., Chang, Y.C., Lewis, A.L., Nizet, V., and Varki, A. (2009). Molecular mimicry of host sialylated glycans allows a bacterial pathogen to engage neutrophil Siglec-9 and dampen the innate immune response. Blood 113, 3333–3336.

Cauci, S., Monte, R., Driussi, S., Lanzafame, P., and Quadrifoglio, F. (1998). Impairment of the mucosal immune system: IgA and IgM cleavage detected in vaginal washings of a subgroup of patients with bacterial vaginosis. The Journal of infectious diseases 178, 1698–1706.

Cho, I., and Blaser, M.J. (2012). The human microbiome: at the interface of health and disease. Nat Rev Genet 13, 260–270.

Coombs, G.H., and North, M.J. (1983). An Analysis of the Proteinases of Trichomonas-Vaginalis by Polyacrylamide-Gel Electrophoresis. Parasitology 86, 1–6.

Corfield, A.P. (2018). The Interaction of the Gut Microbiota with the Mucus Barrier in Health and Disease in Human. Microorganisms 6.

Cott, C., Thuenauer, R., Landi, A., Kühn, K., Juillot, S., Imberty, A., Madl, J., Eierhoff, T., and Römer, W. (2016). *Pseudomonas aeruginosa* lectin LecB inhibits tissue repair processes by triggering beta-catenin degradation. BBA -Molecular Cell Research 1863, 1106–1118.

Cross, B.W., and Ruhl, S. (2018). Glycan recognition at the saliva - oral microbiome interface. Cell Immunol 333, 19–33.

Dawson, N.L., Sillitoe, I., Lees, J.G., Lam, S.D., and Orengo, C.A. (2017). CATH-Gene3D: Generation of the resource and its use in obtaining structural and functional annotations for protein sequences. Methods Mol Biol 1558, 79–110.

Eddy, S.R. (2004). Where did the BLOSUM62 alignment score matrix come from? Nat Biotechnol 22, 1035–1036.

Edgar, R.C. (2004). MUSCLE: multiple sequence alignment with high accuracy and high throughput. Nucleic Acids Res 32, 1792–1797.

Eierhoff, T., Bastian, B., Thuenauer, R., Madl, J., Audfray, A., Aigal, S., Juillot, S., Rydell, G.E., Müller, S., de Bentzmann, S., et al. (2014). A lipid zipper triggers bacterial invasion. Proc Natl Acad Sci U S A 111, 12895–12900.

Etzold, S., and Juge, N. (2014). Structural insights into bacterial recognition of intestinal mucins. Curr Opin Struct Biol 28, 23–31.

Fazli, M., Almblad, H., Rybtke, M.L., Givskov, M., Eberl, L., and Tolker-Nielsen, T. (2014). Regulation of biofilm formation in Pseudomonas and Burkholderia species. Environ Microbiol 16, 1961–1981.

Fettweis, J.M., Serrano, M.G., Brooks, J.P., Edwards, D.J., Girerd, P.H., Parikh, H.I., Huang, B., Arodz, T.J., Edupuganti, L., Glascock, A.L., et al. (2019). The vaginal microbiome and preterm birth. Nat Med 25, 1012–1021.

Ficko-Blean, E., and Boraston, A.B. (2012). Insights into the recognition of the human glycome by microbial carbohydrate-binding modules. Curr Opin Struct Biol 22, 570–577.

Fujimoto, Z., Tateno, H., and Hirabayashi, J. (2014). Lectin structures: classification based on the 3-D structures. Methods Mol Biol 1200, 579–606.

Gipson, I.K. (2001). Mucins of the human endocervix. Front Biosci 6, D1245–1255.

Iliev, I.D., Funari, V.A., Taylor, K.D., Nguyen, Q., Reyes, C.N., Strom, S.P., Brown, J., Becker, C.A., Fleshner, P.R., Dubinsky, M., et al. (2012). Interactions between commensal fungi and the C-type lectin receptor Dectin-1 influence colitis. Science 336, 1314–1317.

Imberty, A., Mitchell, E.P., and Wimmerová, M. (2005). Structural basis for high affinity glycan recognition by bacterial and fungal lectins. Curr Opin Struct Biol 15, 525–534.

Kilian, M., Reinholdt, J., Lomholt, H., Poulsen, K., and Frandsen, E.V.G. (1996). Biological significance of IgA1 proteases in bacterial colonization and pathogenesis: Critical evaluation of experimental evidence. Apmis 104, 321–338.

Kindinger, L.M., Bennett, P.R., Lee, Y.S., Marchesi, J.R., Smith, A., Cacciatore, S., Holmes, E., Nicholson, J.K., Teoh, T.G., and MacIntyre, D.A. (2017). The interaction between vaginal microbiota, cervical length, and vaginal progesterone treatment for preterm birth risk. Microbiome 5, 6.

Kindinger, L.M., MacIntyre, D.A., Lee, Y.S., Marchesi, J.R., Smith, A., McDonald, J.A., Terzidou, V., Cook, J.R., Lees, C., Israfil-Bayli, F., et al. (2016). Relationship between vaginal microbial dysbiosis, inflammation, and pregnancy outcomes in cervical cerclage. Sci Transl Med 8, 350ra102.

Kitts, P.A., Church, D.M., Thibaud-Nissen, F., Choi, J., Hem, V., Sapojnikov, V., Smith, R.G., Tatusova, T., Xiang, C., Zherikov, A., et al. (2016). Assembly: a resource for assembled genomes at NCBI. Nucleic Acids Res 44, D73–80.

Kline, K.A., Falker, S., Dahlberg, S., Normark, S., and Henriques-Normark, B. (2009). Bacterial adhesins in host-microbe interactions. Cell Host Microbe 5, 580–592.

Lepenies, B., and Lang, R. (2019). Editorial: Lectins and Their Ligands in Shaping Immune Responses. Front Immunol 10, 2379.

Lis, H., and Sharon, N. (1998). Lectins: Carbohydrate-specific proteins that mediate cellular recognition. Chem Rev 98, 637–674.

Lombard, V., Golaconda Ramulu, H., Drula, E., Coutinho, P.M., and Henrissat, B. (2014). The carbohydrate-active enzymes database (CAZy) in 2013. Nucleic Acids Res 42, D490–495.

Loris, R. (2002). Principles of structures of animal and plant lectins. Biochim Biophys Acta 1572, 198–208.

Ma, B., Forney, L.J., and Ravel, J. (2012). Vaginal microbiome: rethinking health and disease. Annu Rev Microbiol 66, 371–389.

MacIntyre, D.A., Sykes, L., and Bennett, P.R. (2017). The human female urogenital microbiome: complexity in normality. Emerging Topics in Life Sciences 1, 363–372.

Macklaim, J.M., Gloor, G.B., Anukam, K.C., Cribby, S., and Reid, G. (2011). At the crossroads of vaginal health and disease, the genome sequence of Lactobacillus iners AB-1. Proc Natl Acad Sci U S A 108 Suppl 1, 4688–4695.

Mariethoz, J., Alocci, D., Gastaldello, A., Horlacher, O., Gasteiger, E., Rojas-Macias, M., Karlsson, N.G., Packer, N., and Lisacek, F. (2018). Glycomics@ExPASy: Bridging the gap. Mol Cell Proteomics.

Mariethoz, J., Khatib, K., Alocci, D., Campbell, M.P., Karlsson, N.G., Packer, N.H., Mullen, E.H., and Lisacek, F. (2016). SugarBindDB, a resource of glycan-mediated host-pathogen interactions. Nucleic Acids Research 44, D1243–1250.

Merritt, E.A., and Hol, W.G.J. (1995). AB5 toxins. Curr Opin Struct Biol 5, 165–171.

Mesnage, S., Dellarole, M., Baxter, N.J., Rouget, J.B., Dimitrov, J.D., Wang, N., Fujimoto, Y., Hounslow, A.M., Lacroix-Desmazes, S., Fukase, K., et al. (2014). Molecular basis for bacterial peptidoglycan recognition by LysM domains. Nat Commun 5, 4269.

Mitra, A., MacIntyre, D.A., Ntritsos, G., Smith, A., Tsilidis, K.K., Marchesi, J.R., Bennett, P.R., Moscicki, A.B., and Kyrgiou, M. (2020). The vaginal microbiota associates with the regression of untreated cervical intraepithelial neoplasia 2 lesions. Nat Commun 11, 1999.

Montanier, C., Flint, J.E., Bolam, D.N., Xie, H., Liu, Z., Rogowski, A., Weiner, D.P., Ratnaparkhe, S., Nurizzo, D., Roberts, S.M., et al. (2010). Circular permutation provides an evolutionary link between two families of calcium-dependent carbohydrate binding modules. J Biol Chem 285, 31742–31754.

Moonens, K., and Remaut, H. (2017). Evolution and structural dynamics of bacterial glycan binding adhesins. Curr Opin Struct Biol 44, 48–58.

Moschioni, M., Pansegrau, W., and Barocchi, M.A. (2010). Adhesion determinants of the Streptococcus species. Microb Biotechnol 3, 370–388.

Neelamegham, S., Aoki-Kinoshita, K., Bolton, E., Frank, M., Lisacek, F., Lutteke, T., O’Boyle, N., Packer, N.H., Stanley, P., Toukach, P., et al. (2019). Updates to the Symbol Nomenclature for Glycans guidelines. Glycobiology 29, 620–624.

Nishiyama, K., Sugiyama, M., and Mukai, T. (2016). Adhesion Properties of Lactic Acid Bacteria on Intestinal Mucin. Microorganisms 4.

Pang, X., Xiao, X., Liu, Y., Zhang, R., Liu, J., Liu, Q., Wang, P., and Cheng, G. (2016). Mosquito C-type lectins maintain gut microbiome homeostasis. Nat Microbiol 1, 16023.

Petrova, M.I., Reid, G., Vaneechoutte, M., and Lebeer, S. (2017). Lactobacillus iners: Friend or Foe? Trends Microbiol 25, 182–191.

Potter, S.C., Luciani, A., Eddy, S.R., Park, Y., Lopez, R., and Finn, R.D. (2018). HMMER web server: 2018 update. Nucleic Acids Res 46, W200–W204.

Rampersaud, R., Planet, P.J., Randis, T.M., Kulkarni, R., Aguilar, J.L., Lehrer, R.I., and Ratner, A.J. (2011). Inerolysin, a cholesterol-dependent cytolysin produced by Lactobacillus iners. J Bacteriol 193, 1034–1041.

Reimers, L.L., Mehta, S.D., Massad, L.S., Burk, R.D., Xie, X., Ravel, J., Cohen, M.H., Palefsky, J.M., Weber, K.M., Xue, X., et al. (2016). The Cervicovaginal Microbiota and Its Associations With Human Papillomavirus Detection in HIV-Infected and HIV-Uninfected Women. The Journal of infectious diseases 214, 1361–1369.

Robertson, J.A., Stemler, M.E., and Stemke, G.W. (1984). Immunoglobulin-a Protease Activity of Ureaplasma-Urealyticum. J Clin Microbiol 19, 255–258.

Sayers, E.W., Agarwala, R., Bolton, E.E., Brister, J.R., Canese, K., Clark, K., Connor, R., Fiorini, N., Funk, K., Hefferon, T., et al. (2019). Database resources of the National Center for Biotechnology Information. Nucleic Acids Res 47, D23–D28.

Sehnal, D., Deshpande, M., Varekova, R.S., Mir, S., Berka, K., Midlik, A., Pravda, L., Velankar, S., and Koca, J. (2017). LiteMol suite: interactive web-based visualization of large-scale macromolecular structure data. Nature Methods 14, 1121–1122.

Tailford, L.E., Crost, E.H., Kavanaugh, D., and Juge, N. (2015). Mucin glycan foraging in the human gut microbiome. Front Genet 6, 81.

Thomas-White, K., Forster, S.C., Kumar, N., Van Kuiken, M., Putonti, C., Stares, M.D., Hilt, E.E., Price, T.K., Wolfe, A.J., and Lawley, T.D. (2018). Culturing of female bladder bacteria reveals an interconnected urogenital microbiota. Nat Commun 9, 1557.

Thornton, D.J., Rousseau, K., and McGuckin, M.A. (2008). Structure and function of the polymeric mucins in airways mucus. Annu Rev Physiol 70, 459–486.

UniProt, C. (2019). UniProt: a worldwide hub of protein knowledge. Nucleic Acids Res 47, D506–D515.

van de Wijgert, J., Verwijs, M.C., Gill, A.C., Borgdorff, H., van der Veer, C., and Mayaud, P. (2020). Pathobionts in the Vaginal Microbiota: Individual Participant Data Meta-Analysis of Three Sequencing Studies. Front Cell Infect Microbiol 10, 129.

van de Wijgert, J.H., Borgdorff, H., Verhelst, R., Crucitti, T., Francis, S., Verstraelen, H., and Jespers, V. (2014). The vaginal microbiota: what have we learned after a decade of molecular characterization? PLoS One 9, e105998.

Vaneechoutte, M. (2017). The human vaginal microbial community. Res Microbiol 168, 811–825.

Vasta, G.R., Amzel, L.M., Bianchet, M.A., Cammarata, M., Feng, C., and Saito, K. (2017). F-Type Lectins: A Highly Diversified Family of Fucose-Binding Proteins with a Unique Sequence Motif and Structural Fold, Involved in Self/Non-Self-Recognition. Front Immunol 8, 1648.

Velez, M.P., De Keersmaecker, S.C., and Vanderleyden, J. (2007). Adherence factors of Lactobacillus in the human gastrointestinal tract. FEMS Microbiol Lett 276, 140–148.

Vornhagen, J., Quach, P., Boldenow, E., Merillat, S., Whidbey, C., Ngo, L.Y., Adams Waldorf, K.M., and Rajagopal, L. (2016). Bacterial Hyaluronidase Promotes Ascending GBS Infection and Preterm Birth. MBio 7.

Wiggins, R., Hicks, S.J., Soothill, P.W., Millar, M.R., and Corfield, A.P. (2001). Mucinases and sialidases: their role in the pathogenesis of sexually transmitted infections in the female genital tract. Sex Transm Infect 77, 402–408.

Zhang, H., Yohe, T., Huang, L., Entwistle, S., Wu, P., Yang, Z., Busk, P.K., Xu, Y., and Yin, Y. (2018). dbCAN2: a meta server for automated carbohydrate-active enzyme annotation. Nucleic Acids Res 46, W95–W101.

