## Supplemental tables for "Proteome-wide prediction of bacterial carbohydrate-binding proteins as a tool for understanding commensal and pathogen colonisation of the vaginal microbiome"

### S1 Table : List of lectin classes identified from Unilectin3D and used in the classification

| *Lectin class* | *Fold* | *known 3D structure* | *Bacterial structures in UniLectin3D* | *Terminal sugar recognised* | *Bacterial predictions in specified score interval* | | | |
|---|---|---|---|---|---|---|---|---|
| | | | | | *0-0.25* | *0.25-0.5* | *0.5-1* | *Total* |
| ***chi-lectin TCLL*** | a/b barrel / TIM | Plant | None | b-GlcNAc | 6 | 7 | 5 | 18 |
| ***P-domain of calnexin and reticulin*** | a/b hairpin / non-globular proline-rich | Animal | None | a-Glc | 15 | 7 | 0 | 22 |
| ***C-type lectin*** | a/b mixed / C-type lectin-like | Animal | None | a-Fuc, a-Man | 1088 | 2632 | 0 | 3720 |
| ***LysM-like*** | a/b mixed / LysM domain | Fungal | None | b-GlcNAc | 25201 | 4575 | 0 | 29776 |
| ***Ficolin-like*** | a/b mixed with b-sheet / Fibrinogen C-ter like | Animal | None | b-GlcNAc | 1015 | 46 | 25 | 1086 |
| ***MAR Micronemal protein*** | a/b mixed with b-sheet / MAR_domain | Protist | None | a-NeuAc | 0 | 0 | 0 | 0 |
| ***Factor H-binding protein*** | a/b mixed with b-sheet / not classified | Bacterial | 2 | a-Gal | 11 | 92 | 49 | 152 |
| ***HOP-OMP adhesins*** | a/b mixed with b-sheet / not classified | Bacterial | 15 | a-Fuc | 1754 | 1624 | 698 | 4076 |
| ***Mycoplasma adhesins*** | a/b mixed with b-sheet / not classified | Bacterial | | a-NeuAc | 92 | 37 | 26 | 155 |
| ***Cholera toxin like AB5*** | a/b OB-fold | Bacterial | 33 | b-Gal | 3 | 15 | 156 | 174 |
| ***heat labile toxin IIB AB5*** | a/b OB-fold | Bacterial | 5 | a-NeuAc | 1 | 3 | 54 | 58 |
| ***Pertussis toxin AB5*** | a/b OB-fold | Bacterial | 3 | a-NeuAc | 0 | 1 | 28 | 29 |
| ***Shiga toxin AB5*** | a/b OB-fold | Bacterial | 20 | a-Gal | 18 | 102 | 502 | 622 |
| ***Staphylococcal enterotoxin*** | a/b OB-fold | Bacterial | 2 | b-Gal | 276 | 660 | 198 | 1134 |
| ***Staphylococcal Superantigen-Like*** | a/b OB-fold | Bacterial | 6 | a-NeuAc | 523 | 528 | 777 | 1828 |
| ***Thyphoid toxin like AB5*** | a/b OB-fold | Bacterial | 9 | a-NeuAc | 124 | 227 | 906 | 1257 |
| ***Plasmodium Erythrocyte binding antigen*** | ahelix_triplets / Duffy-like | Protist | None | a-NeuAc | 0 | 0 | 0 | 0 |
| ***OAA-like*** | b-barrel | Bacterial | 10 | a-Man | 14 | 18 | 141 | 173 |
| ***P-type lectin*** | b-barrel | Animal | None | a-Man | 0 | 0 | 0 | 0 |
| ***P-type lectin-like*** | b-barrel | Animal | None | a-Man | 0 | 0 | 0 | 0 |
| ***Toxin repetitive domain*** | b-hairpin stack | Bacterial | | a-Gal | 7238 | 1395 | 48 | 8681 |
| ***Acinetobacter phage AP22*** | b-helix | Virus | None | NA | 52 | 30 | 0 | 82 |
| ***E coli bacteriophage*** | b-helix | Virus | None | a-Rha | 33 | 12 | 19 | 64 |
| ***E coli phage CBA120*** | b-helix | Virus | None | a-Glc | 860 | 4 | 2 | 866 |
| ***Salmonella bacteriophage*** | b-helix | Virus | None | a-Rha | 249 | 130 | 62 | 441 |
| ***Shigella phage Sf6*** | b-helix | Virus | None | a-Rha | 425 | 9 | 18 | 452 |
| ***Jacalin-like*** | b-prism I | Mixed | None | a-Man | 858 | 15 | 0 | 873 |
| ***Natterin-like*** | b-prism I | Animal | None | a-Man | 75 | 14 | 0 | 89 |
| ***Oyster lectin*** | b-prism I | Animal | None | a-Man | 68 | 58 | 1 | 127 |
| ***Vibrio b-prism*** | b-prism I | Bacterial | 12 | a-Man | 74 | 146 | 44 | 264 |
| ***Monocot-lectin like*** | b-prism II | Mixed | None | a-Man | 702 | 2955 | 6 | 3663 |
| ***Pyocin*** | b-prism II | Bacterial | 1 | NA | 162 | 679 | 186 | 1027 |
| ***Fungal prism lectins*** | b-prism III | Fungal | None | a-Fuc | 0 | 0 | 53 | 53 |
| ***AAL-like PropLec6A*** | b-propeller | Mixed | None | a-Fuc | 8 | 77 | 81 | 166 |
| ***BPL and CVL like PropLec7C*** | b-propeller | Bacterial | 2 | NA | 126 | 152 | 17 | 295 |
| ***neuraminidase-like*** | b-propeller | Virus | None | a-NeuAc | 5 | 0 | 0 | 5 |
| ***PLL-like PropLec7A*** | b-propeller | Bacterial | 15 | a-Fuc | 1632 | 2071 | 90 | 3793 |
| ***PVL-like PropLec7B*** | b-propeller | Fungal | None | D-GlcNAc | 668 | 417 | 0 | 1085 |
| ***Tachylectin-2-like PropLec5A*** | b-propeller | Animal | None | D-GlcNAc | 59 | 52 | 0 | 111 |
| ***Tectonin PropLec6B*** | b-propeller | Mixed | None | a-Man | 154 | 328 | 3 | 485 |
| ***2 calcium lectin*** | b-sandwich / 2 ca lectin | Bacterial | 62 | a-Fuc | 8 | 143 | 520 | 671 |
| ***ANXUR1 Malectin-like*** | b-sandwich / ConA-like | Animal | None | D-GlcNAc | 10 | 0 | 0 | 10 |
| ***Coronavirus spike protein*** | b-sandwich / ConA-like | Virus | None | a-NeuAc | 4 | 0 | 0 | 4 |
| ***ERGIC-VIP L-type*** | b-sandwich / ConA-like | Mixed | None | a-Man | 100 | 276 | 0 | 376 |
| ***galectin-like*** | b-sandwich / ConA-like | Mixed | None | b-Gal | 33 | 14 | 0 | 47 |
| ***Laminin G-like*** | b-sandwich / ConA-like | Plant | None | b-Xyl | 12 | 0 | 0 | 12 |
| ***L-type legume lectin*** | b-sandwich / ConA-like | Animal | None | aMan, b-Gal | 214 | 904 | 3 | 1121 |
| ***Malectin*** | b-sandwich / ConA-like | Animal | None | a-Glc | 381 | 1058 | 0 | 1439 |
| ***Pentraxin*** | b-sandwich / ConA-like | Animal | None | NA | 56 | 259 | 0 | 315 |
| ***Physarum lectin*** | b-sandwich / ConA-like | Protist | None | NA | 0 | 2 | 0 | 2 |
| ***Rotavirus spike protein*** | b-sandwich / ConA-like | Virus | None | a-NeuAc | 2 | 0 | 0 | 2 |
| ***Rotavirus spike protein P2*** | b-sandwich / ConA-like | Virus | None | a-NeuAc | 0 | 1 | 0 | 1 |
| ***Yeast Emp L-type*** | b-sandwich / ConA-like | Fungal | None | NA | 0 | 0 | 0 | 0 |

| | | | | | | | | |
|---|---|---|---|---|---|---|---|---|
| ***L-rhamnose binding lectin*** | b-sandwich / CUB-like | Animal | None | a-Rha | 7 | 72 | 5 | 84 |
| ***cyanovirin-like*** | b-sandwich / cyanovirin-like | Mixed | None | a-Man | 62 | 228 | 136 | 426 |
| ***Microvirin CVN-like*** | b-sandwich / cyanovirin-like | Bacterial | 2 | a-Man | 49 | 42 | 84 | 175 |
| ***Fungal fruit body lectin*** | b-sandwich / cytolysin-like | Fungal | None | D-GlcNAc | 36 | 0 | 0 | 36 |
| ***Oomycete cytolysin*** | b-sandwich / cytolysin-like | Protist | None | D-GlcN | 46 | 1320 | 174 | 1540 |
| ***1 Calcium lectin*** | b-sandwich / Galactose-binding domain-like | Bacterial | 24 | a-Gal | 7 | 112 | 160 | 279 |
| ***CBM67-like*** | b-sandwich / Galactose-binding domain-like | Fungal | None | a-Gal | 1408 | 46 | 0 | 1454 |
| ***F-type lectin*** | b-sandwich / Galactose-binding domain-like | Mixed | None | a-Fuc | 303 | 3078 | 174 | 3555 |
| ***H-type lectin*** | b-sandwich / Galactose-binding domain-like | Animal | None | a-GalNAc | 15 | 1014 | 3 | 1032 |
| ***Sea_anemon_lectin*** | b-sandwich / Galactose-binding domain-like | Animal | None | a-Gal | 0 | 0 | 0 | 0 |
| ***Ig-like*** | b-sandwich / Ig-like | Animal | None | NA | 2 | 16 | 0 | 18 |
| ***I-type lectin*** | b-sandwich / Ig-like | Fungal | None | a-NeuAc | 144 | 53 | 0 | 197 |
| ***serine-rich repeat adhesin*** | b-sandwich / Ig-like | Bacterial | 10 | a-NeuAc | 733 | 243 | 24 | 1000 |
| ***PA14 RTX*** | b-sandwich / PA14 adhesin | Bacterial | 2 | a-Glc | 1179 | 358 | 237 | 1774 |
| ***PA14 yeast adhesin*** | b-sandwich / PA14 adhesin | Fungal | None | a-Man | 192 | 33 | 0 | 225 |
| ***bacterial adhesin CfaE*** | b-sandwich / pili and adhesins | Bacterial | 1 | NA | 156 | 851 | 22 | 1029 |
| ***bacterial adhesin FaeG*** | b-sandwich / pili and adhesins | Bacterial | 2 | b-Gal | 38 | 277 | 151 | 466 |
| ***bacterial adhesin FedF*** | b-sandwich / pili and adhesins | Bacterial | 3 | NA | 0 | 0 | 28 | 28 |
| ***bacterial adhesin FimH-FlmH*** | b-sandwich / pili and adhesins | Bacterial | 68 | a-Man | 130 | 338 | 451 | 919 |
| ***bacterial adhesin GafD*** | b-sandwich / pili and adhesins | Bacterial | 13 | b-GlcNAc | 3 | 22 | 41 | 66 |
| ***bacterial adhesin PapG*** | b-sandwich / pili and adhesins | Bacterial | 8 | a-Gal | 51 | 63 | 67 | 181 |
| ***bacterial adhesin PsaA*** | b-sandwich / pili and adhesins | Bacterial | 5 | a-Fuc | 2 | 17 | 58 | 77 |
| ***bacterial adhesin UclD*** | b-sandwich / pili and adhesins | Bacterial | 2 | b-Gal | 15 | 26 | 29 | 70 |
| ***TNFa-like*** | b-sandwich / TNF-like | Bacterial | 4 | a-Fuc | 0 | 23 | 125 | 148 |
| ***Bluetongue virus coat protein*** | b-sandwich / viral coat and capsid protein | Virus | None | a-NeuAc | 3 | 0 | 0 | 3 |
| ***Equine_Rhinitis_A_Virus_Coat_Protein*** | b-sandwich / viral coat and capsid protein | Virus | None | a-NeuAc | 0 | 0 | 0 | 0 |
| ***FMDV receptor complex*** | b-sandwich / viral coat and capsid protein | Virus | None | a-Fuc | 16 | 2 | 1 | 19 |
| ***Polyomavirus capsid protein*** | b-sandwich / viral coat and capsid protein | Virus | None | a-NeuAc | 14 | 2 | 0 | 16 |
| ***hemagglutinin-esterase*** | b-sandwich / viral protein domain | Virus | None | a-NeuAc | 1 | 0 | 0 | 1 |
| ***Influenza hemagglutinin*** | b-sandwich / viral protein domain | Virus | None | a-NeuAc | 5 | 23 | 19 | 47 |
| ***Fiber knob*** | b-sandwich / virus globular domain | Virus | None | a-NeuAc | 1 | 1 | 2 | 4 |
| ***Fiber-knob parvovirus*** | b-sandwich / virus globular domain | Virus | None | a-NeuAc | 10 | 4 | 0 | 14 |
| ***Phage binding domain*** | b-sandwich / virus globular domain | Virus | None | b-GlcNAc | 15 | 0 | 0 | 15 |
| ***Turkey siadenovirus A*** | b-sandwich / virus globular domain | Virus | None | a-NeuAc | 119 | 5 | 0 | 124 |
| ***Amaranthin-like*** | b-trefoil | Plant | None | b-Gal | 0 | 0 | 0 | 0 |
| ***Boletus and Laetiporus b-trefoil lectin*** | b-trefoil | Fungal | None | b-Gal | 83 | 16 | 0 | 99 |
| ***Clitocybe lectin-like*** | b-trefoil | Fungal | None | b-Gal | 2 | 8 | 0 | 10 |
| ***Clostridial toxin*** | b-trefoil | Bacterial | 43 | b-Gal | 258 | 45 | 55 | 358 |
| ***Cys-rich man-receptor*** | b-trefoil | Animal | None | b-Gal | 3 | 4 | 0 | 7 |
| ***Earthworm lectin*** | b-trefoil | Animal | None | b-Gal | 212 | 192 | 0 | 404 |
| ***Fungi and Clostridium b-trefoil lectin*** | b-trefoil | Mixed | None | b-Gal | 29 | 11 | 39 | 79 |
| ***Mussel lectin*** | b-trefoil | Animal | None | a-Gal | 9 | 32 | 0 | 41 |
| ***Ricin-like*** | b-trefoil | Mixed | None | b-Gal | 5140 | 9475 | 0 | 14615 |
| ***Sclerotinia lectin like*** | b-trefoil | Fungal | None | b-Gal | 1 | 3 | 0 | 4 |
| ***Trefoil Factor*** | peptide | Animal | None | a-GlcNAc | 4 | 4 | 3 | 11 |
| ***SML2 Micronemal protein*** | small protein / APPLE domain | Animal | None | b-Gal | 77 | 166 | 0 | 243 |
| ***Cyanobacterial scytovirin*** | small protein / disulfide rich | Bacterial | 3 | NA | 2 | 25 | 6 | 33 |
| ***Invertebrate chitin-binding protein*** | small protein / Invertebrate chitin-binding protein | Animal | None | b-GlcNAc | 0 | 21 | 0 | 21 |
| ***Ginkbilobin*** | small protein / Knottin | Plant | None | a-Man | 1 | 1 | 12 | 14 |
| ***Ginkbilobin-like*** | small protein / Knottin | Fungal | None | NA | 0 | 0 | 0 | 0 |
| ***hevein*** | small protein / Knottin | Plant | None | b-GlcNAc | 1 | 19 | 1 | 21 |
| ***Lyophyllum ginkbilobin-like*** | small protein / Knottin | Fungal | None | a-Gal | 2 | 0 | 0 | 2 |
| ***Spider_toxin*** | small protein / Knottin | Animal | None | NA | 0 | 0 | 0 | 0 |

### S2 Table. List of the species and strains used in the study

| Species and strain | Symptom | Source |
|---|---|---|
| Escherichia_coli_UMB0731 | Pathobiont | PMID 29674608 |
| Escherichia_coli_UMB0789 | Pathobiont | PMID 29674608 |
| Escherichia_coli_UMB0900 | Pathobiont | PMID 29674608 |
| Escherichia_coli_UMB0901 | Pathobiont | PMID 29674608 |
| Escherichia_coli_UMB6721 | Pathobiont | Bioproject PRJNA316969 |
| Escherichia_coli_UMB7431 | Pathobiont | Bioproject PRJNA316969 |
| Gardnerella_vaginalis_DSM4944 | Pathobiont | Feizi Ten Array |
| Gardnerella_vaginalis_GED7275B | Pathobiont | PMID 30633889 |
| Gardnerella_vaginalis_GED7760B | Pathobiont | PMID 30633889 |
| Gardnerella_vaginalis_UMB0032A | Pathobiont | PMID 29674608 |
| Gardnerella_vaginalis_UMB0032B | Pathobiont | PMID 29674608 |
| Gardnerella_vaginalis_UMB0061 | Pathobiont | PMID 29674608 |
| Gardnerella_vaginalis_UMB0170 | Pathobiont | Bioproject PRJNA316969 |
| Gardnerella_vaginalis_UMB0233 | Pathobiont | PMID 29674608 |
| Gardnerella_vaginalis_UMB0264 | Pathobiont | Bioproject PRJNA316969 |
| Gardnerella_vaginalis_UMB0298 | Pathobiont | PMID 29674608 |
| Gardnerella_vaginalis_UMB0386 | Pathobiont | PMID 29674608 |
| Gardnerella_vaginalis_UMB0682 | Pathobiont | PMID 29674608 |
| Gardnerella_vaginalis_UMB0768 | Pathobiont | Bioproject PRJNA316969 |
| Gardnerella_vaginalis_UMB0770 | Pathobiont | PMID 29674608 |
| Gardnerella_vaginalis_UMB0775 | Pathobiont | PMID 29674608 |
| Gardnerella_vaginalis_UMB0830 | Pathobiont | PMID 29674608 |
| Gardnerella_vaginalis_UMB0833 | Pathobiont | PMID 29674608 |
| Gardnerella_vaginalis_UMB0912 | Pathobiont | PMID 29674608 |
| Gardnerella_vaginalis_UMB0913 | Pathobiont | PMID 29674608 |
| Lactobacillus_crispatus_C037 | Commensal | Bioproject PRJNA316969 |
| Lactobacillus_crispatus_MV-1A-US | Commensal | PMID 30633889 |
| Lactobacillus_crispatus_SJ-3C-US | Commensal | PMID 30633889 |
| Lactobacillus_crispatus_UMB0040 | Commensal | Bioproject PRJNA316969 |
| Lactobacillus_crispatus_UMB0044 | Commensal | Bioproject PRJNA316969 |
| Lactobacillus_crispatus_UMB0054 | Commensal | PMID 29674608 |
| Lactobacillus_crispatus_UMB0085 | Commensal | PMID 29674608 |
| Lactobacillus_crispatus_UMB0803 | Commensal | PMID 29674608 |
| Lactobacillus_crispatus_UMB0824 | Commensal | PMID 29674608 |
| Lactobacillus_crispatus_UMB1398 | Commensal | PMID 29674608 |
| Lactobacillus_crispatus_VMC3 | Commensal | PMID 30633889 |
| Lactobacillus_crispatus_VMC4 | Commensal | PMID 30633889 |
| Lactobacillus_crispatus_VMC5 | Commensal | PMID 30633889 |
| Lactobacillus_crispatus_VMC7 | Commensal | PMID 30633889 |
| Lactobacillus_crispatus_VMC8 | Commensal | PMID 30633889 |
| Lactobacillus_delbrueckii_UMB0003 | Commensal | PMID 29674608 |
| Lactobacillus_fermentum_UMB0187 | Commensal | PMID 29674608 |
| Lactobacillus_gasseri_202-4 | Commensal | PMID 30633889 |
| Lactobacillus_gasseri_SJ-9E-US | Commensal | PMID 30633889 |
| Lactobacillus_gasseri_SV-16A-US | Commensal | PMID 30633889 |
| Lactobacillus_gasseri_UMB0045 | Commensal | PMID 29674608 |
| Lactobacillus_gasseri_UMB0045b | Commensal | PMID 29674608 |
| Lactobacillus_gasseri_UMB0099 | Commensal | PMID 29674608 |
| Lactobacillus_gasseri_UMB1399 | Commensal | Bioproject PRJNA316969 |
| Lactobacillus_iners_ATCC55195 | Pathobiont | PMID 30633889 |
| Lactobacillus_iners_DSM13335 | Pathobiont | Feizi Ten Array |
| Lactobacillus_iners_LEAF2052A-d | Pathobiont | PMID 30633889 |
| Lactobacillus_iners_SPIN2503V10-d | Pathobiont | PMID 30633889 |
| Lactobacillus_iners_UMB0030 | Pathobiont | Bioproject PRJNA316969 |
| Lactobacillus_iners_UMB0033 | Pathobiont | Bioproject PRJNA316969 |
| Lactobacillus_iners_UMB1051 | Pathobiont | Bioproject PRJNA316969 |
| Lactobacillus_jensenii_115-3-CHN | Commensal | PMID 30633889 |
| Lactobacillus_jensenii_269-3 | Commensal | PMID 30633889 |
| Lactobacillus_jensenii_SJ-7A-US | Commensal | PMID 30633889 |
| Lactobacillus_jensenii_UMB0007 | Commensal | PMID 29674608 |
| Lactobacillus_jensenii_UMB0034 | Commensal | Bioproject PRJNA316969 |
| Lactobacillus_jensenii_UMB0037 | Commensal | Bioproject PRJNA316969 |
| Lactobacillus_jensenii_UMB0077 | Commensal | PMID 29674608 |
| Lactobacillus_jensenii_UMB1307 | Commensal | Bioproject PRJNA316969 |
| Lactobacillus_jensenii_UMB1355 | Commensal | Bioproject PRJNA316969 |
| Lactobacillus_rhamnosus_51B | Commensal | PMID 30633889 |

| | | |
|---|---|---|
| Lactobacillus_vaginalis_ATCC49540 | Commensal | PMID 30633889 |
| Prevotella_amnii_DNF00058 | Pathobiont | PMID 30633889 |
| Prevotella_amnii_DNF00307 | Pathobiont | PMID 30633889 |
| Prevotella_bivia_DNF00188 | Pathobiont | PMID 30633889 |
| Prevotella_bivia_DNF00320 | Pathobiont | PMID 30633889 |
| Prevotella_bivia_DNF00650 | Pathobiont | PMID 30633889 |
| Prevotella_bivia_GED7760C | Pathobiont | PMID 30633889 |
| Prevotella_corporis_MJR7716 | Pathobiont | PMID 30633889 |
| Prevotella_denticola_DNF00960 | Pathobiont | PMID 30633889 |
| Prevotella_disiens_DNF00882 | Pathobiont | PMID 30633889 |
| Prevotella_timonensis_S9-PR14 | Pathobiont | PMID 30633889 |
| Streptococcus_agalactiae_UMB0049 | Pathobiont | Bioproject PRJNA316969 |
| Streptococcus_agalactiae_UMB0767 | Pathobiont | Bioproject PRJNA316969 |
| Streptococcus_agalactiae_UMB0776 | Pathobiont | Bioproject PRJNA316969 |
| Streptococcus_anginosus_UMB0252 | Pathobiont | PMID 29674608 |
| Streptococcus_anginosus_UMB0820 | Pathobiont | PMID 29674608 |
| Streptococcus_anginosus_UMB0839 | Pathobiont | PMID 29674608 |
| Streptococcus_sanguinis_SK36 | Pathobiont | PMID: 31142849 |
| Streptococcus_mitis_KCOM1350 | Pathobiont | PMID: 31142849 |
| Streptococcus_mitis_CMW7705B | Pathobiont | PMID: 30633889 |
| Streptococcus_mitis_UMB0079 | Pathobiont | PMID 29674608 |
| Streptococcus_mitis_UMB1341 | Pathobiont | PMID 29674608 |
| Streptococcus_parasanguinis_UMB0216 | Pathobiont | PMID 29674608 |
| Streptococcus_salivarius_UMB0051 | Pathobiont | PMID 29674608 |

### S3 Table. CBMs of interest for the present study with associated glycan specificity

| | found in | binding specificity | monosacc | fold |
|---|---|---|---|---|
| CBM3 | bacterial enzymes | mainly cellulose, rarely chitin | b-Glc | β-sandwich |
| CBM4 | bacterial enzymes | xylan, β-1,3-glucan, β-1,3-1,4-glucan, β-1,6-glucan, amorphous but not crystalline cellulose | b-Glc | β-sandwich |
| CBM5 | bacterial enzymes | chitin | b-GlcNAc | |
| CBM6 | bacteria, fungi, horseshoe crab | mostly amorphous cellulose and β-1,4-xylan, rarely β-1,3-glucan, β-1,3-1,4-glucan, and β-1,4-glucan. | b-Glc | β-sandwich |
| CBM8 | bacteria +dictyostelium | cellulose | b-Glc | |
| CBM9 | bacterial xylanases | xylan, rarely cellulose | b-Xyl | β-sandwich |
| CBM11 | bacteria | β-1,4-glucan and β-1,3-1,4-mixed linked glucans | b-Glc | |
| CBM12 | bacteria, fungi | chitin | b-GlcNAc | |
| CBM13 | all | xylan, Man, Gal, GalNAc | b-Xyl | β-trefoil |
| CBM15 | bacteria | xylan, xylo-oligosaccharides | b-Xyl | β-sandwich |
| CBM16 | bacteria | cellulose, glucomannan | b-Glc | β-sandwich |
| CBM17 | bacteria | amorphous and derivatized cellulose, cello-oligosaccharides | b-Glc | |
| CBM20 | all | starch | a-Glc | |
| CBM22 | all | mainly xylan, rarely mixed β-1,3/β-1,4-glucans | b-Xyl | β-sandwich |
| CBM25 | bacteria | starch | a-Glc | β-sandwich |
| CBM26 | bacteria, fungi | starch | a-Glc | β-sandwich |
| CBM27 | bacteria | mannan | b-Man | |
| CBM32 | all | Gal, lactose, LacNAc, polygalacturonic acid | b-Gal | β-sandwich |
| CBM34 | bacteria | granular starch | | |
| CBM35 | all | xylan, mannan, β-galactan, manno-oligosaccharides | b-Xyl | |
| CBM36 | | calcium-dependent binding of xylan, xylo-oligosaccharides | b-Xyl | β-sandwich |
| CBM37 | bacteria | xylan, chitin, microcrystalline and phosphoric-acid swollen cellulose | b-Xyl | |
| CBM38 | bacteria, fungi | inulin | b-Glc | |
| CBM40 | bacteria, worm | sialic acid | a-NeuAc | |
| CBM41 | bacteria, plants | α-glucans amylose, amylopectin, pullulan, and oligosaccharide fragments derived from these polysaccharides | a-Glc | |
| CBM42 | bacteria, fungi | arabinofuranose | a-Ara | β-trefoil |
| CBM45 | plants | starch | a-Glc | |
| CBM46 | bacteria | cellulose | b-Glc | |
| CBM47 | all | fucose | a-Fuc | β-sandwich |
| CBM48 | all | glycogen | a-Glc | |
| CBM49 | plants | crystalline cellulose | b-Glc | |
| CBM51 | bacteria | Gal, blood group A/B-antigens | a-Gal | β-sandwich |
| CBM56 | bacteria, virus | β-1,3-glucan | b-Glc | |
| CBM60 | bacteria | xylan | b-Xyl | |
| CBM61 | bacteria | β-1,4-galactan | b-Gal | |
| CBM63 | bacteria, fungi | cellulose | b-Glc | |
| CBM64 | bacteria | cellulose | b-Glc | |
| CBM65 | bacteria | range of β-glucans/significant preference for xyloglucan | b-Glc | |
| CBM66 | bacteria, fungi | fructans | a-Fru | |
| CBM67 | bacteria, fungi | rhamnose | a-Rha | |
| CBM68 | bacteria | maltotriose | a-Glc | |
| CBM70 | bacteria | hyaluronan | b-GlcA | β-sandwich |
| CBM71 | bacteria | lactose, LacNAc | b-Gal | |
| CBM72 | bacteria | insoluble cellulose, β-1,3/1,4-mixed linked glucans, xylan, and β-mannan | b-Glc | |
| CBM73 | bacteria | chitin | b-GlcNAc | |
| CBM76 | bacteria | xyloglucan, glucomannan, barley β-glucan | b-Xyl | |
| CBM83 | bacteria | starch | a-Glc | |
